# A long read mapping method for highly repetitive reference sequences

**DOI:** 10.1101/2020.11.01.363887

**Authors:** Chirag Jain, Arang Rhie, Nancy Hansen, Sergey Koren, Adam M. Phillippy

**Affiliations:** Department of Computational and Data Sciences, Indian Institute of Science, Bangalore KA 560012, India; Genome Informatics Section, National Human Genome Research Institute, Bethesda, MD 20892, USA, {rhiea, nhansen, sergey.koren, adam.phillippy}@nih.gov

## Abstract

About 5-10% of the human genome remains inaccessible for functional analysis due to the presence of repetitive sequences such as segmental duplications and tandem repeat arrays. To enable high-quality resequencing of personal genomes, it is crucial to support end-to-end genome variant discovery using repeat-aware read mapping methods. In this study, we highlight the fact that existing long read mappers often yield incorrect alignments and variant calls within long, near-identical repeats, as they remain vulnerable to *allelic bias*. In the presence of a non-reference allele within a repeat, a read sampled from that region could be mapped to an incorrect repeat copy because the standard pairwise sequence alignment scoring system penalizes true variants.

To address the above problem, we propose a novel, long read mapping method that addresses allelic bias by making use of *minimal confidently alignable substrings* (MCASs). MCASs are formulated as minimal length substrings of a read that have unique alignments to a reference locus with sufficient mapping confidence (i.e., a mapping quality score above a user-specified threshold). This approach treats each read mapping as a collection of confident sub-alignments, which is more tolerant of structural variation and more sensitive to paralog-specific variants (PSVs) within repeats. We mathematically define MCASs and discuss an exact algorithm as well as a practical heuristic to compute them. The proposed method, referred to as Winnowmap2, is evaluated using simulated as well as real long read benchmarks using the recently completed gapless assemblies of human chromosomes X and 8 as a reference. We show that Winnowmap2 successfully addresses the issue of allelic bias, enabling more accurate downstream variant calls in repetitive sequences. As an example, using simulated PacBio HiFi reads and structural variants in chromosome 8, Winnowmap2 alignments achieved the lowest false-negative and false-positive rates (1.89%, 1.89%) for calling structural variants within near-identical repeats compared to minimap2 (39.62%, 5.88%) and NGMLR (56.60%, 36.11%) respectively.

Winnowmap2 code is accessible at https://github.com/marbl/Winnowmap

## 1 Introduction

Advances in single-molecule sequencing technologies have inspired community efforts to produce high-quality human genome assemblies with accurate resolution of repetitive DNA. The complete, gapless, telomere-to-telomere (T2T) assembly of a human chromosome X is a recent breakthrough that involved assembling a 3.1 Mbp long centromeric satellite DNA array [1]. Similarly, a T2T assembly of human chromosome 8 spanned a 2.1 Mbp long centromere and the 0.6 Mbp long defensin gene cluster for the first time [2]. Such developments are steering genomics into an exciting new era where repeats that were previously thought intractable (e.g., segmental duplications, satellite, and ribosomal DNAs) will no longer remain out of reach. Genomic variants, expected to be in higher abundance within and near repetitive DNA, may be contributing to complex traits such as lifespan and disease [3–5]. PacBio and Oxford Nanopore (ONT) sequencing, due to their orders of magnitude longer read lengths than Illumina, can easily span many common duplications (e.g., LINEs) in the human genome. However, accurate long read mapping within > 100 kbp-sized repeats remains challenging.

Prior algorithmic developments for long read mapping have been crucial to resolving many repetitive sequences and complex variants. As such, several specialized methods have been published to improve long-read match seeding and extension [6–17]. The extension stage typically involves the optimization of a base-to-base alignment score which rewards matching bases while appropriately penalizing gaps and mismatches. However, these alignment scores do not always favor the correct loci in long near-identical repeats, because reads that include non-reference alleles will be penalized and their true loci may score worse than other copies of the repeat. Occurrence of this allelic bias (a.k.a. reference bias) and its effect on estimates of variation and allele frequencies has been extensively discussed in the literature [18–23]. An analogous problem also occurs during genome assembly validation and polishing when reads are mapped back to a potentially erroneous draft assembly [1, 24].

Compared to point mutations or short indels, structural variants (SVs) affect more bases in the genome due to their larger size, and therefore, are bigger contributors to allelic bias. Most existing solutions to address this bias involve modifying the reference sequence, e.g., by adopting a graph-based representation which incorporates known genomic variation [20, 25–27]. While this remains a promising and complementary direction, here we seek to address allelic bias by developing a new long read mapping method that is robust to the presence of novel variation. In our proposed method, referred to as Winnowmap2, we introduce the concept of minimal confidently alignable substrings (MCASs), which are minimal-length read substrings that align end-to-end to a reference with mapping quality score above a user-specified threshold. Through MCASs, we can identify the correct mapping target of a read by considering the substrings that *do not* overlap non-reference alleles. In theory, the mapping quality of each substring quantifies the probability that it is correctly placed [28]. This framework draws advantage from paralog-specific variants (PSVs) [29, 30], which allow us to differentiate repeat copies from each other and identify the MCAS. We provide a formal definition of MCAS, an exact dynamic programming algorithm to compute them, as well as fast heuristics to scale this method to large mammalian genomes.

Winnowmap2 was empirically validated using both simulated and real human genome sequencing benchmarks. In both cases, we judge Winnowmap2 along with the currently available long read mappers by the downstream accuracy of SV calls produced by the SV caller Sniffles [10]. The simulation uses SURVIVOR’s SV benchmarking tool [31] which mutates a reference sequence (in our case, the first two completed human chromosomes 8 and X). Winnowmap2 alignments consistently enabled the most accurate SV calls using both ONT and PacBio HiFi data at varying coverage levels, when compared to other commonly used long read mappers. Using simulated HiFi reads from chromosome 8 (146 Mbp) sampled at 40x coverage, Winnowmap2, minimap2 and ngmlr achieved accuracy scores, i.e., false-negative and false-positive rates (FNR, FPR) of (0.09%, 0.18%), (3.36%, 0.93%) and (3.64%, 2.93%) respectively. Winnowmap2’s improved handling of allelic bias was particularly evident within 7 Mbp of the most repetitive regions of chromosome 8, achieving (FNR, FPR) scores of (1.89%, 1.89%) in these regions compared to (39.62%, 5.88%) for minimap2 and (56.60%, 36.11%) for ngmlr, respectively. Winnowmap2 also achieved favorable accuracy on the Genome in a Bottle (GIAB) benchmark set [32], which excludes long > 10 kbp-sized repeats of the human genome.

## 2 Results

### An overview of the Winnowmap2 algorithm

If an error-free read is simulated directly from a reference, then its correct mapping to that reference computed using a reasonable pairwise sequence alignment algorithm is naturally guaranteed to have the highest score. However, this guarantee does not hold if the same read is mapped to an alternate reference. Consequently, using a pairwise sequence alignment scoring system to judge the *best* mapping candidate is sub-optimal, and this is particularly true while mapping reads to highly repetitive sequences. Regardless of the type of scoring function used, e.g., with either a linear or an affine gap penalty, the function would also penalize variant-induced differences between the sequenced individual and the reference sequence. In cases where one of the repeat copies in a reference sequence contains a different allele from the sequenced individual, reads may achieve a better alignment score against an incorrect repeat copy (Figure 1). An ideal scoring system should ignore non-reference bases when computing an optimal alignment, but these are typically unknown *a priori*.

**Fig. 1.**
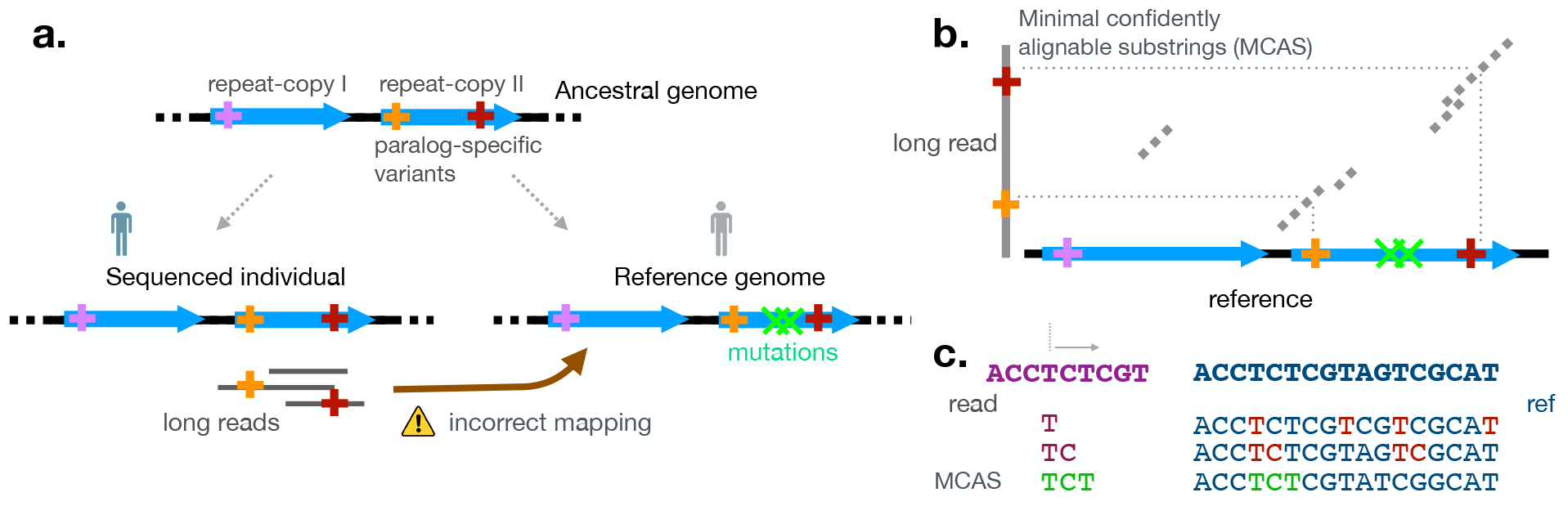
**a.** Illustration of allelic bias in near-identical genomic repeats. Paralog-specific variants (PSVs), indicated using colored ‘+’ markers, are the variants that are unique to a specific repeat copy in an ancestral human genome. Mutations in the reference sequence are indicated using ‘x’ markers. Long reads can be mapped to an incorrect repeat copy if the best mapping is decided by pairwise sequence alignment score. **b.** MCAS alignments map to correct loci on the reference. An MCAS is a carefully selected substring of a read. By excluding non-reference alleles, this approach reduces allelic bias. **c.** MCAS computation is shown using a simple toy example. Starting from several positions in a read, we identify minimum-length substrings that can be uniquely mapped to a reference. Uniqueness of an alignment is determined by using its mapping quality score.

Like most read mappers, Winnowmap2 follows a seed-and-extend workflow. The seeding step reuses Winnowmap’s weighted minimizer sampling [14], which yields an accuracy improvement over the standard minimizer technique [33]. Winnowmap2’s extend stage introduces a novel heuristic to tackle allelic bias. We split the extend stage into two steps. The first step involves identifying minimal confidently alignable substrings (MCASs) from each read to a reference. *Informally, an MCAS at position i of a read refers to the minimum length read substring starting at the position i that achieves a ‘unique’ end-to-end alignment to a reference locus* (see Methods for a formal definition). Here, uniqueness of an alignment is evaluated using a mapping quality (mapQ) score [28] that reflects the score gap between the best and second-best alignment candidates for a substring. Accordingly, an MCAS is valid if its alignment achieves a mapQ score above a user-specified threshold. A read can have as many MCASs as its length. By using MCAS alignments, read bases on either side of a variant can map uniquely to their correct reference loci as they can be scored independently from non-reference bases (Figure 1).

Starting from any position of a read, minimal length of the MCAS is ensured by iteratively increasing substring length and checking whether its maximum scoring alignment to a reference satisfies the mapQ cutoff. Suppose a read is sampled from a repetitive region, the frequency of PSVs at its correct mapping loci helps determine the length of an MCAS. The higher the number of PSVs, the smaller the length of the MCAS, because its mapQ cutoff will be satisfied at an earlier iteration with fewer aligned bases. Similarly, better raw read accuracy also leads to shorter MCAS lengths, since more PSVs will be matched by a more accurate sequence. Shorter MCAS lengths help not only in terms of the runtime with fewer iterations spent but also in terms of accuracy, as MCASs are less likely to overlap non-reference bases.

Computing all MCAS alignments from a read in an exact manner can be computationally prohibitive (see Methods for complexity analysis). To bypass this issue, we rely on banded-alignment and mapQ scoring heuristics from minimap2 [12] to compute each MCAS. For the sake of efficiency, we avoid evaluating MCAS alignments from each consecutive position in a read. Rather, we identify MCAS alignments from a subset of positions that are equally spaced (e.g., 500 bp apart).

The final step in Winnowmap2 is to consolidate a read’s MCAS alignments into a final alignment output. For various reasons (e.g. sequencing errors, approximation of mapQ computation, and complex sequence variants) some MCASs may be incorrectly mapped. During the consolidation step, we extract anchors that participate in each MCAS alignment, and re-execute the chaining and alignment extension algorithm by using the complete set of anchors to output a final alignment. A few anchors from false MCAS mappings are filtered out during the anchor chaining process. We will empirically show that the proposed strategy improves mapping accuracy in repetitive DNA while remaining highly scalable.

### Advantage of Winnowmap2 illustrated using the β-defensin gene cluster

We visualize the advantage of Winnowmap2 method by using the beta-defensin gene cluster on human chromosome 8 as an example. The 7 Mbp beta-defensin locus (chr8: 6,300,000-13,300,000) of the human genome is known to be a hotspot of copy-number variation [34]. In the sequenced CHM13 human cell line, this locus spans three large (> 500 kbp) segmental duplications [2]. To evaluate long read mapping accuracy at this locus, we simulated ONT reads from chr8 at 40x sequencing coverage by using NanoSim [35] (Methods). In addition, we artificially mutated chr8 by adding a 1 kbp deletion variant at position 12,000,000. This locus was chosen for our illustration as it overlaps with one of the three duplications. If mapped correctly, the 1 kbp simulated deletion in the reference should appear as a 1 kbp-long insertion in the overlapping read alignments.

Figure 2 shows an IGV visualization of primary alignments computed by Winnowmap2 and three other long read mapping tools NGMLR, minimap2 and graphmap. Among the four methods, Winnowmap2 achieved the expected mapping coverage in this region with most read alignments showing the expected insertion call. The other tools mapped fewer reads successfully, resulting in reduced coverage and poor read mapQ scores. When these alignments were used as input to Sniffles, only Winnowmap2 alignments resulted in the true SV call. NGMLR, minimap2 and graphmap rely on pairwise sequence alignment scores across the full length of the read when choosing the best mapping target. Due to the large deletion penalty levied at the mutated (but correct) locus, the majority of reads were incorrectly mapped to the other two duplications. Among the three methods, NGMLR showed the least bias, but most of its correct alignments were associated with poor mapQ scores (< 10). A low mapQ score indicates a marginal alignment score difference between the best and the second-best mapping candidate, and therefore, the read alignment may not be considered by the variant caller. This result illustrates the previously discussed limitation of using pairwise alignment scores to rank candidate alignments in genomic repeats. The use of MCASs in Winnowmap2 enabled correct read placements in this case. A few MCAS alignments computed by Winnowmap2 in this region are visualized as a dot-plot in Supplementary Figure S1.

**Fig. 2.**
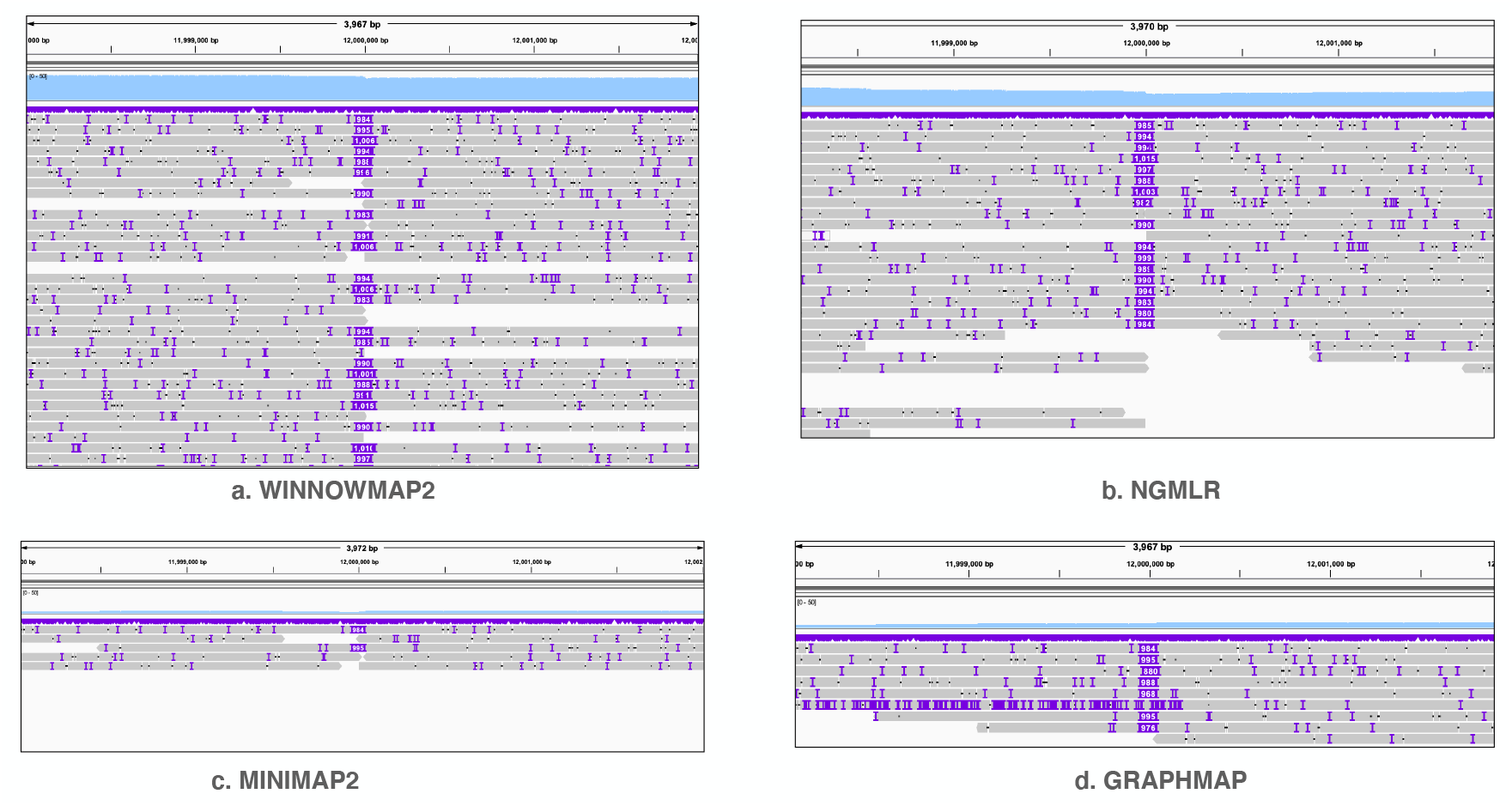
Visualization of alignment pileup near the mutated bases of chromosome 8 by using IGV tool [36]. The sky-blue-colored track on top of each plot shows mapping-coverage using a uniform y-axis scale (0-50). The grey-colored line segments show individual primary read alignments. IGV uses purple markers to indicate presence of indels within read alignments. NGMLR, minimap2, graphmap show reduced coverage due to allelic bias whereas Winnowmap2 shows expected coverage in this region. Consistent large insertions in the middle of each plot are distinctly visible due to simulated SV.

### Evaluation using a simulated benchmark and T2T human chromosomes

We simulated long reads, both HiFi (using PBSIM [37]) and ONT (using NanoSim [35]), at coverage levels of 20x and 40x from T2T assemblies of chromosome 8 (146 Mbp) and chromosome X (154 Mbp) respectively (Methods). To evaluate how well Winnowmap2 addressed allelic bias, we also simulated 1100 structural variants, including both indels (1000) and inversions (100) of size ≤ 1 kbp, in each reference chromosome sequence by using the SURVIVOR benchmarking tool [31]. Both the SV simulation and evaluation of variant sets against the ground truth were done using SURVIVOR (Methods).

We evaluated Winnowmap2, Winnowmap, minimap2 and NGMLR in this experiment to check their false-negative and false-positive rates (FNR, FPR), as well as runtime and memory requirements. The long read mappers produced SAM-formatted alignments, which were then fed to Sniffles [10] to compute SVs. A false negative indicates that a true SV is not supported by read alignments whereas a false positive indicates that a false SV is supported. As such, these statistics are good indicators of the correctness of read alignments. We also performed a *de novo* repeat annotation of each reference sequence (chr8 and chrX) by using Mashmap [38] to identify repetitive sequence intervals of length *≥* 10 kbp and identity *≥* 95% (Supplementary Figure S2). The identified repetitive intervals constitute a notable portion of the two chromosomes; 4.8% in chr8 and 6.9% in chrX. This allowed us to separately evaluate the accuracy of read mappers in near-identical repeats where Winnowmap2 is expected to perform better.

Figures 3a, 3b show the accuracy statistics of the four mapping tools. Winnowmap2 achieved the best FNR and FPR for both the HiFi and ONT read sets. Winnowmap2 FNR and FPR scores consistently stayed below 3% and 0.3% respectively. Most of these gains appear in repetitive sequences of the two chromosomes as evident from our accuracy evaluation within only the repetitive regions (Figures 3c, 3d). Winnowmap2 succeeds in addressing allelic bias in these regions by pre-serving good accuracy in complex repeats where the other tools struggle. These gains were made uniformly over all SV types-insertions, deletions and inversions that were simulated (Supplementary Table S1). When increasing coverage from 20x to 40x, FNR generally reduces for all methods as better sensitivity is naturally expected with higher sequencing coverage.

**Fig. 3.**
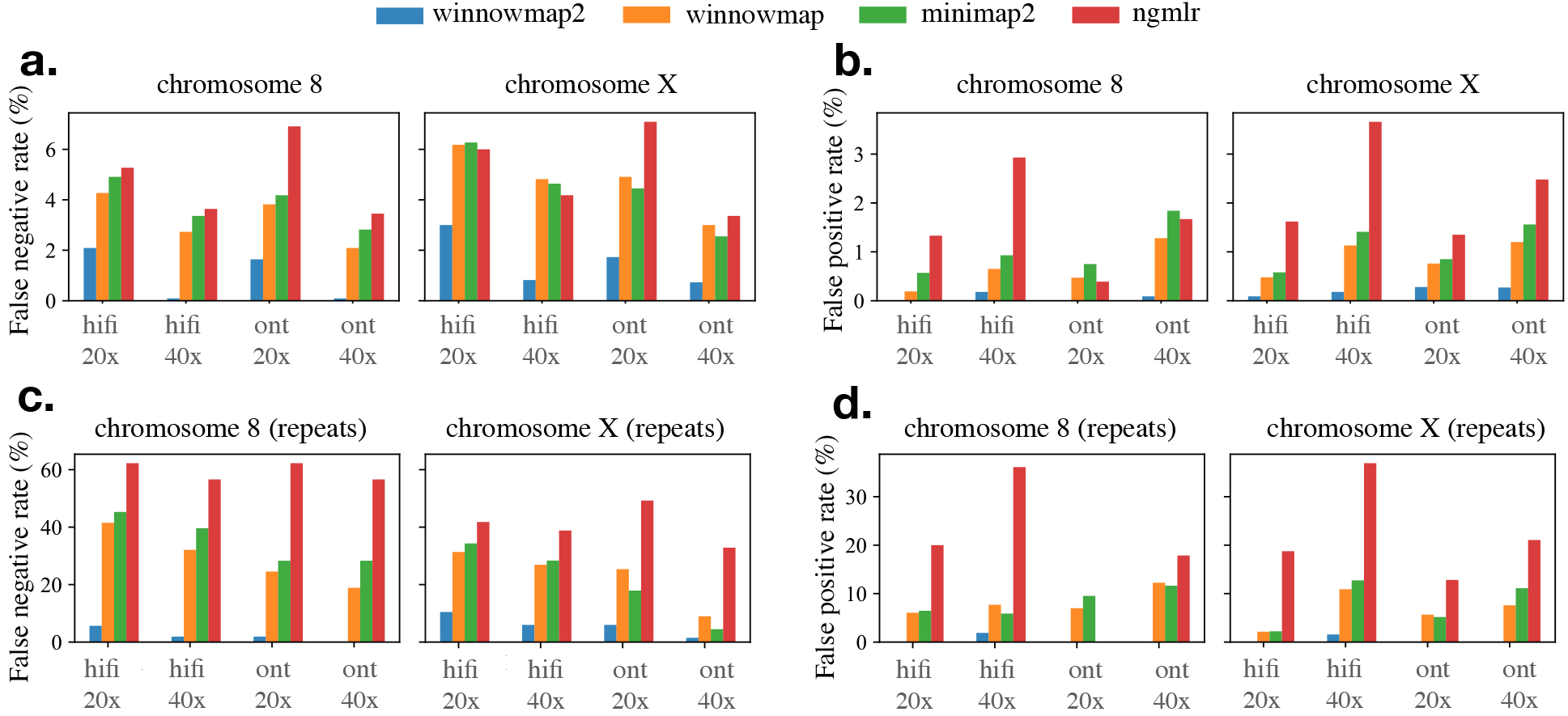
False negative and false positive rates achieved by SV calls of four mapping methods: Winnowmap2, Winnowmap, minimap2 and NGMLR. The top two plots show accuracy statistics over T2T chromosomes 8 and X whereas the bottom two plots show the statistics within only the most repetitive intervals of these chromosomes. Winnowmap2 alignments enabled the most accurate Sniffles SV calls with the least FNR and FPR scores. Note that y-axis scales differ in these plots.

The Winnowmap2 implementation is optimized to run fast while using less memory (Figure 4). As several substring alignments need to be identified from a single read, it requires execution of alignment routines several times rather than just once. In this experiment, minimap2 consistently used the least time, but Winnowmap2 is competitive as its runtime was consistently lower than NGMLR and roughly double minimap2’s runtime.

**Fig. 4.**
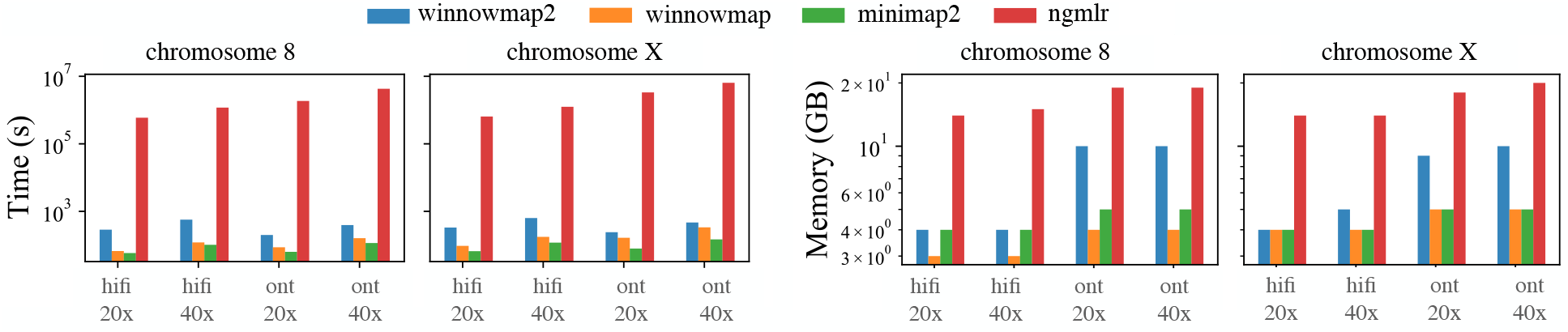
Wall-clock time and memory usage of four mapping methods. Winnowmap2 uses competitive resources when compared to competing methods. Each method was executed using 28 threads on an Intel Xeon processor with 28 physical cores.

### Evaluation using Genome in a Bottle benchmark

Evaluating mappers on real sequencing data is challenging without a known truth. The Genome in a Bottle (GIAB) Tier1 v0.6 benchmark set provides a high-quality characterization of SVs in the Ashkenazi cell line HG002 relative to the GRCh37 human reference. This call set encompasses 2.51 Gbp of the genome and includes 5262 insertions and 4095 deletions [32]. It excludes SVs overlapping segmental duplications and tandem repeats greater than 10 kbp. Nevertheless, this experiment was useful to validate that Winnowmap2 also achieves good mapping accuracy on real data within the commonly studied regions of the genome. Here we mapped three publicly available HG002 long read sequencing sets: HiFi (14-15 kbp library, 35x), ONT (Guppy 3.6.0, 35x) and ONT (Guppy 3.6.0, 50x) to GRCh37, and compared results with minimap2. Similar to our simulated benchmark, variants were called using Sniffles. Winnowmap2 achieved slightly better precision and similar recall scores compared to minimap2 (Figure 5), with similar runtime and memory requirements. We also observed that both Winnowmap2 and minimap2 achieved better SV accuracy using ONT data over HiFi with equal 35x coverage.

**Fig. 5.**
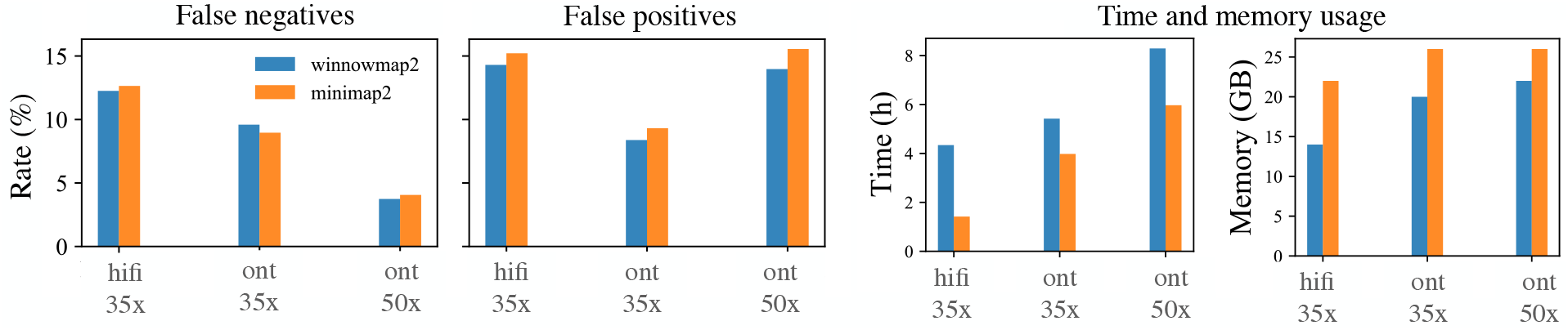
Comparison of Winnowmap2 and minimap2 by using GIAB SV benchmark set defined for HG002 human sample. Current GIAB benchmark set (v0.6) excludes complex repeats of the human genome. Outside the repeats, Winnowmap2 achieves similar FNR scores and slightly better FPR scores compared to minimap2.

## 3 Discussion and Conclusions

Availability of long-range sequencing technologies makes it feasible to resolve large mega-base sized near-identical duplications in the human genome, a feat that was impossible to achieve using short reads alone [39–43]. These regions include recently diverged segmental duplications, ampliconic gene arrays, rRNA genes, and centromeres, all of which play important functional roles in the genome and all of which go largely unstudied by current variation analyses. As human reference gaps associated with these regions are progressively being resolved, this opens up the opportunity to expand the resolution of resequencing approaches. In this work, we highlighted that allelic bias becomes a major challenge for accurately mapping reads to repetitive reference sequences. This challenge affects the accuracy of existing mappers because classic pairwise sequence alignment scoring schemes are not an ideal mechanism to identify the correct mapping target in a repetitive sequence. In Winnowmap2, we have implemented a new idea based on minimal confidently alignable substrings that can be mapped independently of non-reference bases, thus alleviating allelic mapping bias.

We highlighted the advantages of Winnowmap2 by demonstrating its superior downstream variant call accuracy compared to commonly used long read mappers. In particular, Winnowmap2 enabled notable gains in SV calling accuracy within the repetitive regions of human chromosomes. Although here we focused on structural variants, it is natural to expect that Winnowmap2’s superior mapping accuracy will also benefit SNP and short indel variant calling. Prior studies have suggested high enrichment of SVs in repetitive regions that currently correspond to unresolved gaps in the human genome reference [3, 44, 5]. This underscores the importance of understanding how these regions differ between individuals.

Further algorithmic improvements will be needed to improve read alignment accuracy. In particular, it remains challenging to align bases precisely when multiple SVs are clustered in close vicinity. Our simulation made use of SURVIVOR, which simulates SVs at uniformly random positions in a reference sequence and could be an over-simplification of real data. In addition, read mappers and variant callers still remain limited in their ability to handle nested variation and other forms of complex rearrangements [45].

## 4 Methods

### Minimal confidently alignable substring (MCAS)

MCASs distinguish Winnowmap2 from previous read mapping methods. Prior to defining an MCAS, we formalize when we can confidently say that an alignment of a substring is correct. In practice, this confidence is derived using the score difference between the best scoring alignment and other candidate alignments. The mapping quality (mapQ) score was originally defined to address this problem [28], but the existing mathematical definition is restricted to short reads because alignments were assumed to be ungapped. However, the majority of long read sequencing errors are indels and the longer reads are more likely to span structural variants. When allowing for indels, adjacent mapping loci in a reference can no longer be considered independent, as in prior models. Accordingly, we propose the following formulation.

Given a query string *S* and a reference *R*, the top scoring end-to-end (a.k.a. semi-global) alignment candidates of string *S* to *R* can be directly computed in *O*(|*S*| · |*R*|) time. From a pairwise alignment of *S* to *R*, we can identify the set of *matched base positions* between them. For instance, this set would include a tuple (*i, j*) if character *S*[*i*] is matched to character *R*[*j*]. We say that two alignment candidates do not *overlap* if and only if their corresponding sets are disjoint. We associate our confidence with the best-scoring alignment of string S to reference R if and only if its second-best non-overlapping alignment candidate has a score < τ *·* opt, where opt refers to the optimal alignment score and τ ∈ (0, 1) is a user-specified parameter.

Let *Q* be a long read sequence. *A minimal confidently alignable substring MCAS*(*i*) *of read Q refers to the shortest substring starting at position i that has a confident end-to-end alignment to reference R*. For a given read *Q*, we seek MCAS(*i*) ∀ 0 ≤ i < |*Q*|, and their corresponding alignments. MCASs can have variable lengths and can overlap one another. Note that existence of MCAS(*i*) depends on whether it is possible to satisfy the confidence criteria. In the worst case where two repetitive regions lack any PSV (i.e., 100% identical duplicates), then a read sampled from either repeat copy will not contain an MCAS. The rationale of introducing the MCAS idea is to address allelic bias; whereas a non-reference SV allele will cause mis-alignment in the traditional approach, the MCASs are treated independently and those neighboring the SV will remain unaffected (Figure 6).

**Fig. 6.**
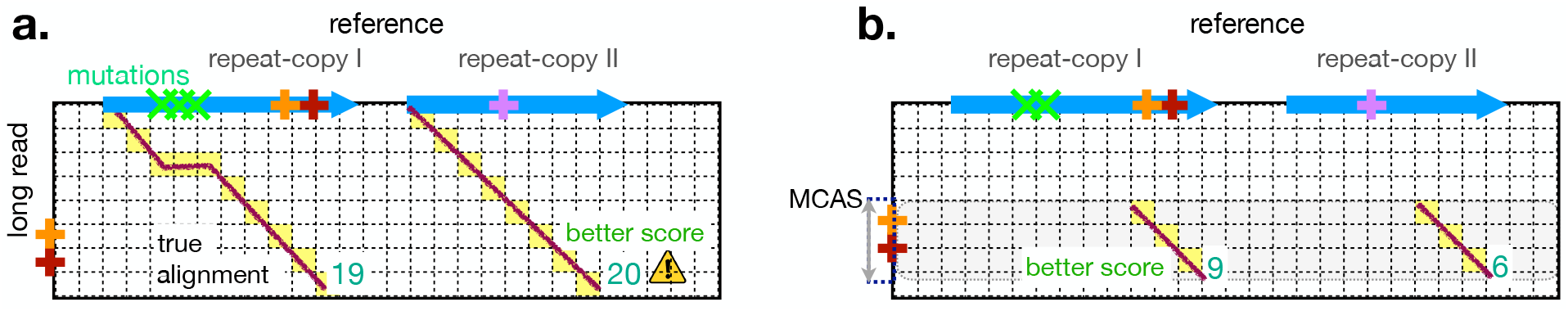
Illustration of MCASs using a DP alignment scoring matrix. Similar to Figure 1, PSVs are shown using colored ‘+’ markers. The left figure highlights the effect of allelic-bias on read alignment scores. The score of a true alignment spanning non-reference alleles can be lower compared to the score of an incorrect alignment. On the right side, an MCAS of a read which does not span non-reference alleles can achieve correct and unique placement to a reference.

Considering the issue of allelic bias, it is also desirable to enforce a maximum length parameter for valid MCASs because long MCASs again become vulnerable to allelic bias. By default, we set the maximum length parameter to 8 kbp for HiFi reads and 16 kbp for ONT reads based on our experimental observations. As such, the maximum length of a valid MCAS is a constant. The lemma below summarizes the asymptotic complexity to compute MCASs.

#### Lemma 1.

*Computing MCAS(i*) ∀ 0 ≤ i < |*Q*| *requires O*(|*Q*||*R*|) *time and O(|R|) space*.

*Proof.* Assume any appropriate linear or affine gap scoring function is being used. Denote *p*^th^ character in read Q as Q[*p*] and a substring ranging from positions *p* to *q* as *Q[p, q]* with both ends inclusive. Consider the following algorithm to compute MCAS(*i*). Compute semi-global DP alignment of *Q*[*i*, *i+j*] to reference *R* while iterating the variable *j* from 0 to *c*. Here *c* is the maximum length allowed for a valid MCAS. Computing a row of the alignment scoring table requires *O(|R|)* time. As a new row is computed in an iteration, we need to check whether the best-scoring alignment satisfies the confidence criteria, i.e., whether its score compared to the second-best non-overlapping alignment exceeds by a user-specified threshold. To check this, we require the following additional steps.

Select the cell in the current row associated with the highest score. Next, we need to locate the maximum score achieved by a non-overlapping alignment. Assuming gap penalty is positive, then there can be at most O(j) cells in the DP row that are associated with overlapping alignments. As a result, if we consider cells in non-increasing order of alignment scores, we may need to check at most *O(j)* other cells. *O(j)* cells with the highest scores can be selected from the complete row in *O(|R|)* time using a partition-based selection algorithm. Assume that we also have scores of previous rows in memory. Then we can trivially check for overlap between a pair of alignments in O(j) time. Adding all the work, total asymptotic time spent per row is *O*(*|R| +j*^2^). As total number of rows that may need to be computed is bounded by the constant c, total time spent to compute MCAS(i) remains *O(|R|)*. Therefore, computing all MCASs requires *O(|Q||R|)* time. Asymptotic space complexity of the above algorithm is *O(|R|)*.

An *O(|Q||R|)* time complexity resembles the complexity of DP-based alignment algorithms. As such, the exact algorithm does not offer desired scalability. In Winnowmap2, we make use of fast heuristics and make careful accuracy-performance trade-offs to address this. First, we perform the MCAS computation from a subset of equally spaced starting positions, e.g., after every 500^th^ base. Next, while computing an MCAS, we reduce the alignment search space by making use of known minimizer seeding and clustering ideas [12, 14]. Starting from a small substring length, our iterative method exponentially grows the substring (rather than growing linearly). In each iteration, we check its mapping to reference R. This is done until the substring either satisfies the alignment confidence criteria or cannot be extended further. While computing each mapping, we rely on efficiently engineered anchor chaining, banded-alignment, and mapQ computation code from minimap2. In a way, the mapQ scoring heuristic in minimap2 approximates our definition for confidence assessment.

### Heuristic to compute mapping quality

In Winnowmap2 implementation, we use the same heuristic as minimap2 to compute the mapping quality score of a read alignment. For completeness, we also mention it here. Once the anchors between a read and a reference are identified, minimap2 runs a co-linear chaining algorithm to locate alignment candidates. The chaining procedure ensures that alignment candidates use a disjoint set of anchors to prevent overlaps. To compute mapQ, minimap2 compares the anchor chaining score of the best-scoring chain relative to the second-best. Suppose their scores are denoted as *f*_1_ and *f*_2_ respectively. Also, let m be the count of anchors chained along the best alignment. Minimap2 uses the following empirical formula to calculate mapQ score of the best alignment candidate:

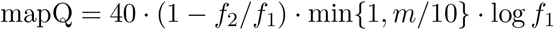

The above score is readjusted by minimap2 to fall within the range of 0 to 60. By default, we use mapQ cutoff of 5 in Winnowmap2 to mark an MCAS alignment as confident. This cutoff can be modified by users. In practice, a higher cutoff typically leads to longer MCASs, as expected. A lower cutoff increases the probability of an incorrect alignment to be considered as the best.

### Consolidating several MCASs into a single alignment output

Once we compute MCAS alignments from a read, these need to be aggregated into a single alignment output. At this step, we extract anchors that were joined to form each MCAS alignment. Subsequently, the union of all anchor sets is passed to chaining and alignment routines to output the final set of best-scoring alignments. Typically, there are only a few anchors to process at this step, which does not require significant time.

### Simulation and evaluation of structural variant calls

In our simulation benchmark, we made use of T2T chromosome assemblies for chromosome 8 (v9) and chromosome X (v0.7) that are available from https://github.com/nanopore-wgs-consortium/CHM13. SURVIVOR (v1.0.6) was used to simulate 1,100 SVs of length ranging from 50 bp to 1000 bp in each chromosome sequence. We also simulated PacBio HiFi reads as well as ONT reads using PBSIM (commit:e014b1) and NanoSim (v2.6.0) respectively. Command line parameters provided to these tools are listed in Supplementary Table S2. NanoSim requires real data for training its error model. Training was executed using a publicly available R10.3 Guppy 3.4.5 ONT sequencing data of the *Escherichia coli* K12 genome (ENA:PRJEB36648). PBSIM command line parameters were adjusted to achieve PacBio HiFi data characteristics with an indel error rate of about 1%. Supplementary Table S3 specifies the read length statistics. Long read mappers were tested using two sequencing coverage levels, 20x and 40x. In our mapping evaluation, we compared Winnowmap2 (v2.0), Winnowmap (v1.01), minimap2 (v2.17), ngmlr (v0.2.7) and graphmap (v0.5.2). Each mapper was executed using their recommended parameters and 28 CPU threads (Table S2). SV calling from BAM alignment file outputs was done using Sniffles (v1.0.11). The SV call sets were evaluated using SURVIVOR against its own simulated ground truth. We also evaluated SV calling accuracy within repetitive reference intervals. For this, *de novo* repeat annotation of reference sequences was computed by using Mashmap (commit:ffeef4) to approximately identify all duplications of *≥*10 kbp length and *≥* 95% identity. SV evaluation within the repeats was done by intersecting variant coordinates and repeat intervals using bedtools (v2.29.2) [46].

### Evaluation using GIAB SV calls

We evaluated Winnowmap2 and minimap2 using the GIAB Tier1 (v0.6) SV call set [32] available for the HG002 human sample relative to the GRCh37 human genome reference. In this experiment, we utilized HG002 ONT and PacBio HiFi sequencing data [47, 48] made available through the precision FDA site https://precision.fda.gov/challenges/10/. Sniffles SV call sets were evaluated using SVanalyzer (v0.36).

## Acknowledgements

We would like to thank Kishwar Shafin, Alla Mikheenko, and Sergey Nurk for providing useful feedback regarding Winnowmap2. We also acknowledge Heng Li for responding to our queries regarding minimap2 code. Winnowmap2 and Winnowmap were developed on top of minimap2 code. This research was supported in part by the Intramural Research Program of the National Human Genome Research Institute, National Institutes of Health.

