## Supplementary Material for "A long read mapping method for highly repetitive reference sequences"

---

### Content summary

- **Figure S1:** Visualisation of MCAS read alignments using a simulated ONT read and chr8 reference sequence.
- **Figure S2:** Annotation of long near-identical duplications in CHM13 chromosome 8 and X respectively.
- **Table S1:** Structural variant accuracy evaluation using chromosome 8 and chromosome X as reference sequences respectively.
- **Table S2:** Command line parameters that were used to execute various tools for this study.
- **Table S3:** Length statistics of simulated and real long read sequencing data sets used in this study.

---

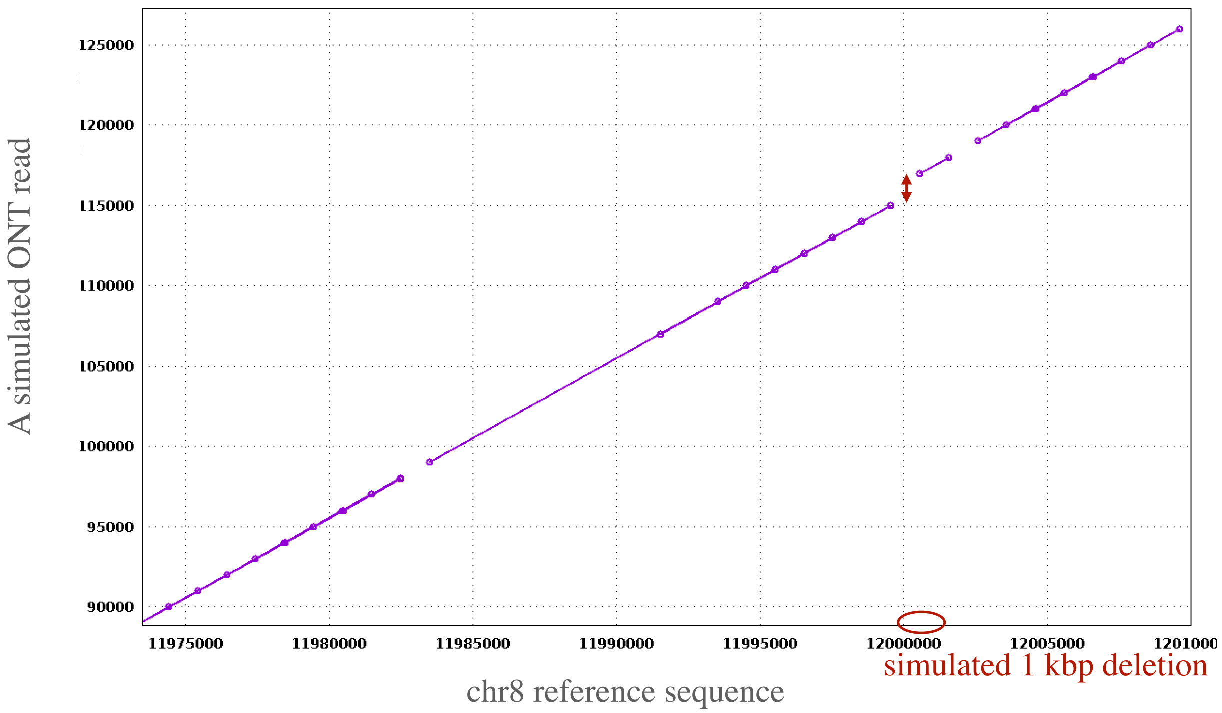


**Fig. S1.** Several MCAS alignments computed by Winnowmap2 are shown for a simulated ONT read using a dot-plot. MCASs surrounding the non-reference SV allele are correctly aligned to the mutated chr8 reference sequence. A purple dot indicates either the start or the end of an MCAS alignment. MCASs can have variable length and can also overlap with each other. These MCAS alignments are joined together in a final step by Winnowmap2.

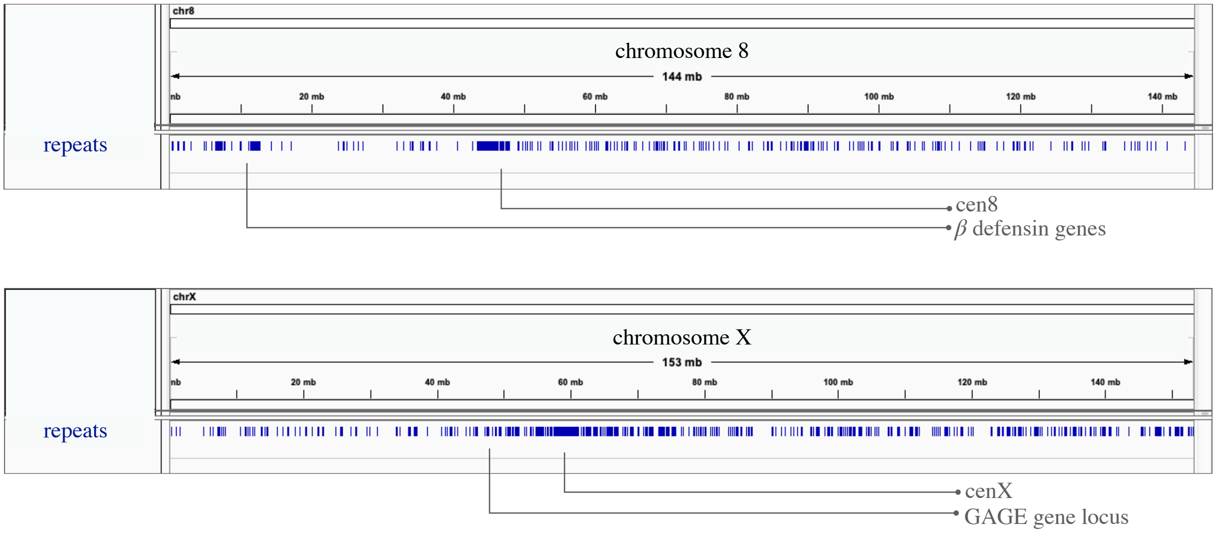


**Fig. S2.** De novo annotation of long near-identical duplications in CHM13 chromosome 8 and X respectively. In each case, these duplications (length$\geq$ 10 kbp, identity$\geq$ 95%) were identified by executing self-alignment of these chromosomes using Mashmap. Mashmap (https://github.com/marbl/MashMap) includes a script that generates these intervals in bed format. A few repeat units are also labeled in the above figure.

**chromosome 8**, total calls simulated 510/100/490

| Dataset | | Winnowmap2 | Winnowmap | minimap2 | MGMLR |
|---|---|---|---|---|---|
| hifi 20x | Total calls | 500/98/479 | 494/85/474 | 492/85/469 | 493/95/454 |
| | FN calls | 10/2/11 | 16/15/16 | 18/15/21 | 17/5/36 |
| | FP calls | 0/0/0 | 1/0/1 | 3/0/3 | 6/2/0 |
| hifi 40x | Total calls | 509/100/490 | 503/87/480 | 501/87/475 | 497/97/466 |
| | FN calls | 1/0/0 | 7/13/10 | 9/13/15 | 13/3/24 |
| | FP calls | 1/0/1 | 5/0/2 | 6/0/4 | 12/4/0 |
| ONT 20x | Total calls | 501/97/484 | 494/86/478 | 493/85/476 | 479/93/452 |
| | FN calls | 9/3/6 | 16/14/12 | 17/15/14 | 31/7/38 |
| | FP calls | 0/0/0 | 3/0/2 | 4/0/4 | 2/0/0 |
| ONT 40x | Total calls | 510/100/489 | 505/88/484 | 502/88/479 | 497/97/468 |
| | FN calls | 0/0/1 | 5/12/6 | 8/12/11 | 13/3/22 |
| | FP calls | 0/0/1 | 7/1/6 | 8/2/9 | 6/3/0 |

**chromosome X**, total calls simulated 488/100/512

| Dataset | | Winnowmap2 | Winnowmap | minimap2 | MGMLR |
|---|---|---|---|---|---|
| hifi 20x | Total calls | 476/98/493 | 470/81/481 | 469/80/482 | 468/96/470 |
| | FN calls | 12/2/19 | 18/19/31 | 19/20/30 | 20/4/42 |
| | FP calls | 1/0/0 | 4/1/0 | 4/1/1 | 6/6/0 |
| hifi 40x | Total calls | 485/98/508 | 473/81/493 | 472/81/496 | 471/97/486 |
| | FN calls | 3/2/4 | 15/19/19 | 16/19/16 | 17/3/26 |
| | FP calls | 2/0/0 | 8/2/0 | 9/3/3 | 14//12/1 |
| ONT 20x | Total calls | 476/99/506 | 467/83/496 | 467/84/500 | 457/90/475 |
| | FN calls | 12/1/6 | 21/17/16 | 21/16/12 | 31/10/37 |
| | FP calls | 2/0/1 | 6/1/1 | 7/1/1 | 5/3/1 |
| ONT 40x | Total calls | 484/99/509 | 480/84/503 | 481/84/507 | 472/97/494 |
| | FN calls | 4/1/3 | 8/16/9 | 7/16/5 | 16/3/18 |
| | FP calls | 2/0/1 | 8/2/2 | 11/3/2 | 8/9/1 |

**Table S1.** Structural variant accuracy evaluation using chromosome 8 and chromosome X as reference sequences respectively. In this experiment, the following three types of SVs were simulated using SURVIVOR: deletions, inversions and insertions. Accordingly all figures of the form $x/y/z$ indicate $x$ deletions, $y$ inversions and $z$ insertions respectively. This table provides a detailed breakdown of the plot shown in main text (Figure 3).

| Tool | Purpose | Command line parameters |
|---|---|---|
| Winnowmap2 (v2.0) | HiFi read mapping | `-W repetitive_k15.txt -ax map-pb --MD ref.fasta hifi.fq.gz` |
| | ONT read mapping | `-W repetitive_k15.txt -ax map-ont --MD ref.fasta ont.fq.gz` |
| minimap2 (v2.17) | HiFi read mapping | `-t 28 -ax asm20 --MD ref.fasta hifi.fq.gz` |
| | ONT read mapping | `-t 28 -ax map-ont --MD ref.fasta ont.fq.gz` |
| ngmlr (v0.2.7) | HiFi read mapping | `-t 28 -r ref.fasta -q hifi.fq.gz -o output.sam` |
| | ONT read mapping | `-t 28 -x ont -r ref.fasta -q ont.fq.gz -o output.sam` |
| Winnowmap (v1.01) | HiFi read mapping | `-W repetitive_k19.txt -t 28 -ax asm20 --MD ref.fasta hifi.fq.gz` |
| | ONT read mapping | `-W repetitive_k15.txt -t 28 -ax map-ont --MD ref.fasta ont.fq.gz` |
| graphmap (v0.5.2) | ONT read mapping | `align -r ref.fasta -d ont.fq.gz -o output.sam -t 28` |
| Sniffles (v1.0.11) | SV calling | `-n -1 -t 8 -m alignments.bam -v output.vcf` |
| SURVIVOR (v1.0.6) | SV simulation | `simSV ref.fasta parameter_file 0 1 alternate` |
| | SV evaluation | `eval SV.vcf truth.bed 50 results` |
| SVanalyzer(v0.36) | SV evaluation | `SVbenchmark --ref ref.fasta --test test.vcf --truth truth.vcf --includebed giab.bed --testfilter PASS --truthfilter PASS --normdist 1.00 --normsizediff 1.00 --normshift 1.00` |
| PBSIM (commit:e014b1) | HiFi read simulation | `--depth 20 --data-type CLR --accuracy-mean 0.999 --accuracy-min 0.99 --length-min 18000 --length-mean 20000 --length-max 22000 --model_qc model_qc_clr ref.fasta` |
| NanoSim(v2.6.0) | ONT read simulation | `read_analysis.py genome -i ont.fq.gz -rg ref1.fasta -ga output.sam -t 28 -o train, simulator.py genome -rg ref2.fasta -c train -med 50000 -sd 0.5 -t 28 -n $NUM` |
| Mashmap (commit:ffeef4) | Repeat annotation | `mashmap -r ref.fasta -q ref.fasta -f none -s 10000 --pi 95, python denovo_repeat_annotation.py mashmap.out 10000 95 > tmp.bed, bedtools merge -i tmp.bed` |
| bedtools(v2.29.2) | Filter SVs within repeats | `bedtools intersect -a output.vcf -b repeats.bed -u -wa > repeats.vcf` |

**Table S2.** Command line parameters that were used to execute various tools for this study.

Simulated long reads from T2T chromosomes 8 and X that were used in this study

| Dataset | Read count | N50 | Min. length | Max. Length | Notes |
|---|---|---|---|---|---|
| HiFi (chr8, 20x) | 146,118 | 19,979 | 18,000 | 22,000 | simulated by PBSIM |
| HiFi (chr8, 40x) | 292,286 | 19,972 | 18,000 | 22,000 | simulated by PBSIM |
| HiFi (chrX, 20x) | 154,869 | 19,969 | 18,000 | 22,000 | simulated by PBSIM |
| HiFi (chrX, 40x) | 309,745 | 19,971 | 18,000 | 22,000 | simulated by PBSIM |
| ONT (chr8, 20x) | 51,993 | 63,283 | 8,915 | 369,183 | simulated by NanoSim |
| ONT (chr8, 40x) | 103,723 | 63,503 | 4,498 | 497,739 | simulated by NanoSim |
| ONT (chrX, 20x) | 54,073 | 64,260 | 11,076 | 331,049 | simulated by NanoSim |
| ONT (chrX, 40x) | 110,227 | 63,763 | 1,272 | 342,759 | simulated by NanoSim |

Real human HG002 sequencing data used in this study

| Dataset | Read count | N50 | Min. length | Max. Length | Notes |
|---|---|---|---|---|---|
| HiFi (HG002, 35x) | 8,449,287 | 12,885 | 47 | 30,581 | 15 kbp library |
| ONT (HG002, 35x) | 12,563,983 | 50,382 | 1 | 543,308 | Guppy 3.6.0 PromethION |
| ONT (HG002, 50x) | 19,328,993 | 50,380 | 1 | 543,308 | Guppy 3.6.0 PromethION |

**Table S3.** Length statistics of simulated and real long read sequencing data sets used in this study.
